# Estimation of population divergence times from SNP data and a test for treeness

**DOI:** 10.1101/281881

**Authors:** Christoph Theunert, Montgomery Slatkin

## Abstract

We present a method for estimating population divergence times from genome sequences when one individual is sampled from each population. Our method is a simplified version of one presented by Rassmussen et al. (2014) for testing for direct ancestry of an archaic genome. Our method does not require distinguishing ancestral from derived alleles or assumptions about demographic history before population divergence. We discuss the relationship of our method to two similar methods, one introduced by Green et al. (2010) and denoted by *F*(*A* | *B*) and the other introduced by Schlebusch et al. (2017) and called the TT method. When our method is applied to individuals from three or more populations, it provides a test of whether the population history is treelike. We illustrate the use of our method by applying it to three high-coverage archaic genomes, two Neanderthals (Vindija and Altai) and a Denisovan.

---

Several methods have been proposed for estimating the divergence time of populations under the assumption that they have remained isolated since they diverged. For distantly related populations, the numbers of mutational differences between sequences indicates relative times of divergence. Relative times can be converted to absolute times if the mutation rate is known. This method traces to Zuckerkandl and Pauling (1962; 1965) and has been used and refined extensively. This class of methods estimates genomic divergence times. Using it to estimate population or species divergence times assumes that divergence times are so large that the difference between genomic and population divergence can be ignored.

For recently diverged populations, the number of mutational differences probably does not provide a reliable estimate of population divergence times both because there may be too few mutations that distinguish populations and because the difference between the genomic and population divergence times may be substantial. To overcome this problem, Green et al. (2010) (in Supplement 14) introduced a method for estimating population divergence times that accounts for the difference between genomic and population divergence. This method was used in later papers from the same group (MEYER *et al.* 2012; PRÜFER *et al.* 2014; PRÜFER *et al.* 2017).

The Green et al. (2010) method is applicable when one genome is sampled from each of two populations. It depends on the statistic *F*(*A*|*B*), which is the fraction of sites in population *A* that carry the derived allele when that site is heterozygous in population *B*. Green et al. (2010) showed by simulation that *F*(*A*|*B*) decreases roughly exponentially with the separation time of populations *A* and *B*.

The pattern of decrease depends on the history of population sizes in *B* and in the population ancestral to *A* and *B.*

In a different context, Rasmussen et al. (2014) (Supplement 17) introduced a method for inferring whether an archaic sample was in a population directly ancestral to a present-day population. Although the Rasmussen et al. (2014) method appears to be different from the *F*(*A*|*B*) method, it is actually quite similar. It uses analytic expressions for the number of configurations of pairs of sites in the two populations, when no distinction is made between ancestral and derived alleles. It makes no assumptions about the history of population sizes in either population.

More recently, Schlebusch et al. (2017) in Section 9.1 of their supplementary materials, introduced another and similar method, called the TT method, for estimating population divergence times. Their method is based on analytic expressions for the seven configurations of SNPs that are polymorphic in the two populations. The TT method assumes ancestral and derived alleles can be distinguished and the population before divergence was of constant size.

In the present paper, we will derive a simpler version of the Rassmussen et al. method and describe the relationship to the *F*(*A*|*B*) and TT methods. We will show that our method provides a way to test whether the population history of three or more populations is accurately represented by a population tree.

## Analytic theory of *F*(*A*|*B*)

We assume that two populations *A* and *B* diverged at time *T* in the past and remained isolated since. Two chromosomes are sampled from population *B* and one from *A*. Let *N*(*t*) denote the population size *t* generations before the present (*t*=0). Between 0 and *T, N*(*t*) is the effective size of population *B*. Before *T*, it is the effective size of the ancestral population. Because only one chromosome is sampled from *A*, the effective size of *A* between 0 and *T* does not matter. If there is no recurrent mutation. *A* carries the derived allele only if one of the two B lineages coalesced with the *A* lineage and there was a mutation on the internal branch of the gene tree, as shown in Figure 1. We calculate the probability of those two events using standard coalescent theory.

**Figure 1.**
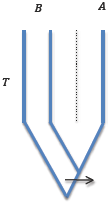
Illustration of the notation used in this paper. Populations *A* and *B* are assumed to have diverged from a common ancestor *T* generations in the past. Two chromosomes from *B* and one from *A* are sampled. A mutation from the ancestral to derived allele at a SNP is assumed to have occurred on the gene tree as shown by the arrow. Assuming no recurrent mutation, the gene tree and the branch on which the mutation occurred is the only way that *B* can be heterozygous for the derived allele and *A* also carry the derived allele.

The probability of the gene tree shown in Figure 1 is 2(1 – *c*) / 3 where *c* is the probability that the two *B* lineages coalesce between 0 and *T*. The 2/3 reflects the fact that in the ancestral population each pair of lineages is equally likely to coalesce first. The probability that there is coalescence between 0 and *T* is

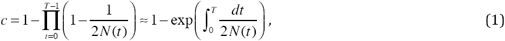

where the approximation is accurate when *N*(*t*) is large.

If *N* is constant,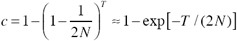.

We denote the expected length of the internal branch of the gene tree shown in Figure 1 by *u*. In general *u* depends on *N*(*t*) in a complicated way but if *N* is constant, *u* = 2*N* (WAKELEY 2009). The probability that a mutation occurs in the internal branch is *µu* where *µ* is the per site mutation rate.

The unconditional probability that the two *B* lineages carry different alleles is *µ* multiplied by twice the average coalescence time of those two lineages, 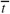, where

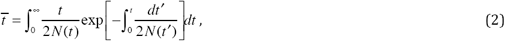

Note that 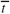 does not depend on *T*. When *N* is constant 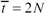.

We denote the probability that *A* carries the derived allele given that the two *B* lineages carry different alleles by *P*(*A* | *B*). We distinguish this probability from the statistic *F*(*A* | *B*) computed from the data. From the rules of conditional probability we obtain

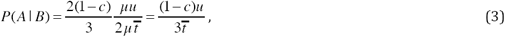

which reduces to *P*(*A* | *B*) = exp[–*T* / (2*N*)] / 3 when *N* is constant. Note that in Equation (3) the mutation rate cancels. If *N* varies with time, an analytic expression for *P*(*A* | *B*) can be obtained for some functional forms of *N*(*t*), but in practice it is easier to determine the dependence of *P*(*A* | *B*) on *T* by simulation, as was done by Green et al. (2010) and in later papers.

Green et al. (2010) estimated the decrease in *P*(*A* | *B*) with time for several demographic models and then estimated *T* by finding the intersection point with the observed value of *F*(*A* | *B*) with each simulated curve. In Equation (3), *P*(*A* | *B*) depends on *N*(*t*) both before and after *T* because *u* depends on *N*(*t*) before and after *T*.

## Schlebusch et al. (2017) TT method

A closely related method for estimating population divergence times was presented by Schlebusch et al. (2017) in part 9 of their supplemental materials (pp. 21-23). They call this method the TT method and note that it is closely related to the concordance methods previously used by Schlebusch et al. (2012) and Skoglund et al. (2011). Schlebusch et al. (2017) assume that two chromosomes are sampled from each population and distinguish nine configurations of the data at each site: O0 (0/0), O1 (1/0), O2 (0/1), O3 (2/0), O4 (0/2), O5 (1/1) O6 (2/1), O7 (1/2) and O8 (2/2), where the numbers before and after the slash are the numbers of derived alleles in the first and second populations respectively. Schlebusch et al. derived the probabilities of each configuration under the infinite sites model with constant mutation rate, arbitrary population size changes after population separation, and constant population size in the ancestral population before separation. These probabilities depend on several parameters: the probabilities of coalescence in the two daughter populations, here called *c*_1_ and *c*_2_to be consistent with the notation in the previous section, *T*_1_and *T*_2_ (the population split times for each population), *V*_1_and *V*_2_(the expected times to coalescence in the two populations, given that they coalesce before the populations split), and *θ*, (the effective size of the ancestral population scaled by the mutation rate). They assume that the numbers of sites in each of the nine configurations take their expected values for a given sample size, and they derived expressions for each of the parameters. In particular, they showed that the two coalescence probabilities are given by

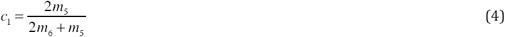

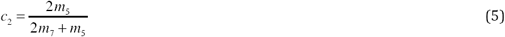

where *m*_*i*_ is the observed number of sites in configuration O*i*.

## Rasmussen et al. method

Rasmussen et al. (2014) (Supplement 17) considered the problem of whether an archaic genome was from a population directly ancestral to a present-day population. Like the TT method, two chromosomes are sampled from each population. The two populations *A* and *B* were assumed to have separated at some time in the past. To eliminate mutation as a force, they restricted their analysis to sites that were ascertained to be polymorphic in an outgroup. We call the two alleles by S and s. Without distinguishing ancestral and derived alleles, there are five different configurations of the data at each site: (1) SS/SS or ss/ss, (2) SS/Ss or ss/Ss, (3) SS/ss or ss/SS, (4) Ss/SS or Ss/ss, and (5) Ss/Ss, where the first genotype is from population *A* and the second is from *B*. Rassmussen et al. (2014) showed that, in the absence of mutations, the probabilities of the five configurations depend on five parameters, *c*_1_, the probability that the two lineages from *A* coalesce after the populations diverge, *c*_2_, the probability that the two lineages from *B* coalesce after the populations diverge, and *k*_0_, *k*_1_and *k*_2_, the elements of the normalized folded site-frequency spectrum in a sample of size 4 immediately before the populations diverged: *k*_0_is the probability of SSSS or ssss, *k*_1_is the probability of SSSs or sSSS, and *k*_2_is the probability of SSss, where the ordering of S and s does not matter.

The data consist of the numbers of sites *n*_i_ with each configuration. Rasmussen et al. (2014) assumed the data had a multinomial distribution with probabilities *p*_*i*_. They used standard numerical methods for estimating the five parameters from the data. This is a composite likelihood method because linkage disequilibrium can cause nearby sites to be non-independent.

Rasmussen et al. (2014) applied their method to an archaic sample from Montana, which in this notation is population *B*, and several present-day Native American individuals, each of which in turn was population *A*. Rasmussen et al. restricted their analysis to sites that are polymorphic in a panel of African populations. As a test of direct ancestry, Rasmussen et al. used a likelihood ratio test of the hypothesis that *c*_2_=0. If *c*_2_=0, the branch to *B* from the population ancestral to *A* and *B* was so short that no coalescence events occurred. In doing this analysis, Rasmussen et al. (2014) needed no assumptions about the history of population sizes either before or after *T*.

In this paper, we simplify the Rasmussen et al. (2014) method slightly and assume only one chromosome is sampled from population *A*, as in the *F*(*A* | *B*) method. Also as with the *F*(*A* | *B*) method, there is no need to assume that *A* is a present-day population or even that it was from a more recent time than *B*. The goal is to estimate the coalescence probability in population *B* before populations *A* and *B* had a common ancestor. From that coalescence probability and assumptions about population size changes in *B*, we can estimate *T*, the time since *B* separated from the common ancestor.

With only one chromosome sampled from *A*, there are three configurations of the data: (1) S/SS or s/ss, (2) S/Ss or s/Ss and (3) S/ss or s/SS, where the allele carried by the chromosome from A is before the slash. There are only three parameters of the model, *c*, the probability of coalescence in *B*, and *k*_0_and *k*_1_, the elements of the normalized folded site frequency spectrum in a sample of size 3 at *T*: *k*_0_is the probability of SSS or sss and *k*_1_is the probability of SSs or Sss. There are only two free parameters because *k*_0_ + *k*_1_ = 1. By analogy with the derivation in Rasmussen et al. (2014) (Supplement 17),

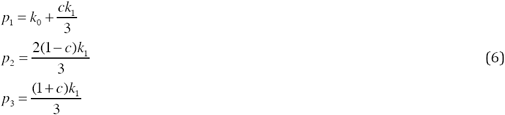

where the *p*_*i*_ are the probabilities of configurations 1, 2 and 3. Given the data, *n*_*i*_ for *i*=1, 2, 3, the three parameters can be estimated by assuming a multivariate normal distribution of the data. From *ĉ*, the estimated value of *c*, the estimated value of *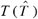* is estimated by solving

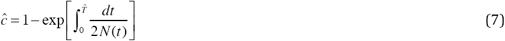

for 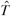. If *N* is constant 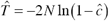We can understand the relationship to *F*(*A*|*B*) by assuming the sample sizes are large enough that the probabilities take their expected values. In that case,

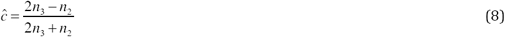

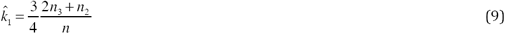

where *n* = *n*_1_ + *n*_2_ + *n*_3_.

The *F*(*A*|*B*) method described in the previous section is similar. To apply it, ancestral and derived alleles must be distinguished. Let S be the derived allele. There are 6 configurations of the data (1) S/SS, (2) S/Ss, (3) S/ss, (4) s/SS, (5) s/Ss and (6) s/ss. Let *v*_*i*_ be the observed numbers of sites in each configuration. By definition,

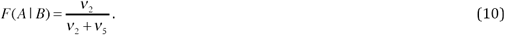

When ancestral and derived alleles are not distinguished, *n*_1_ = *v*1 + *v*_6_, *n*_2_ = *v*_2_ + *v*_5_ and *n*_3_ = *v* _3_ + *v* _4_. Hence, from (8),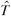 is estimated from

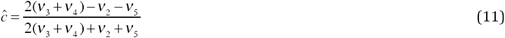

There are several differences between the two methods. First, the two methods use different subsets of the polymorphic sites *F*(*A* | *B*) uses all sites that are heterozygous in *B* while our method uses all sites that are polymorphic in an outgroup. Second, the *F*(*A* | *B*) method estimates *T* directly from simulations, unless *N* is assumed to be constant, while our method first estimates *c* and from that value estimates *T*. Given the assumptions about demography, *c* is an analytic function of *T* which can be solved numerically. No simulations are needed. Third, the estimate of *T* from our method does not depend on the history of population sizes in the common ancestral population. Those population sizes determine *k*_1_ which is estimated from the data.

It is more difficult to compare our method and the *F*(*A* | *B*) method with the TT method because the TT method assumes that two chromosomes are sampled from each population. For that reason, the TT method is more similar to the Rasmussen et al. (2014) method. One difference is that Rasmussen et al. restricted their analysis to sites polymorphic in an outgroup, thereby eliminating the effects of mutation. The TT method analyzes all polymorphic sites and explicitly accounts for mutation. However, the estimated coalescent probabilities do not depend on the mutation rate (Equations 4 and 5 above). Unfortunately, it is not possible to present an algebraic comparison of the Rasmussen et al. and TT methods because the five equations presented by Rasmussen et al. do not appear to have an analytic solution.

## Application to Neanderthals and Denisovans, with a test of treeness

We illustrate the application of our method to three high-coverage archaic genomes, the Altai Neanderthal from the Denisova Cave in central Siberia (PRÜFER *et al.* 2014), the Vindija Neanderthal from the Vindija Cave in Croatia (PRÜFER *et al.* 2017) and the Denisova genome (MEYER *et al.* 2012). All three genomes were sequenced to sufficient depth that heterozygous sites can be called with confidence. We ascertained a set of SNPs that are polymorphic in a panel of 40 African genomes and restricted our analysis to those SNPs. We used an additional filtering step for the Altai genome. Prüfer et al. (2014) showed that the Altai Neanderthal was inbred with an estimated inbreeding coefficient of 1/8. For the comparisons involving this individual, only sites not in runs of homozygosity longer than 2 mb were analyzed.

With three populations, there are six possible comparisons using each population in turn as population *A* or *B*. Table 1 shows the number of sites in each of the three configurations for all combinations. In the table, one of two alleles chosen at random from population *A* and two from population *B* were analyzed. The estimated value of *c* is the probability of coalescence in population *B* after it diverged from population *A*.

**Table 1.** Counts of SNPs in each of the three configurations along with estimates of *k*_1_ and *c*. Both chromosomes are sampled from population *B* and one chromosome chosen at random is sampled from population *A*. Sites were ascertained to be polymorphic in 40 African individuals in the Simons Genome Diversity Panel (BantuHerero, BantuKenya, BantuTswana, Biaka, Dinka, Esan, Gambian, Ju_hoan, Khomani_San, Luhya, Luo, Mandenka, Masai, Mbuti, Mende, Mozabite, Saharawi, Somali, Yoruba) (Malik et al. 2016). *n*_1_, *n*_2_and *n*_3_are the numbers of sites in each of the three configurations defined in the text.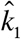 and *ĉ* are obtained by assuming the *n*_*i*_ have a trinomial distribution with probabilities given by Equations (6) in the text and maximizing the likelihood. The same estimates of 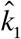 and *ĉ* were obtained using Equations (7) and (8) in the text.

| $B$ | $A$ | $n_1$ | $n_2$ | $n_3$ | $\hat{k}_1$ | $\hat{c}$ |
| --- | --- | --- | --- | --- | --- | --- |
| Altai | Vindija | 7864107 | 75478 | 57311 | 0.017696 | 0.20591 |
| Vindija | Altai | 7874698 | 54704 | 67494 | 0.017790 | 0.42323 |
| Vindija | Denisova | 8980266 | 52108 | 553786 | 0.090730 | 0.91013 |
| Denisova | Vindija | 8973604 | 64912 | 547644 | 0.090771 | 0.88810 |
| Altai | Denisova | 7923816 | 78910 | 465399 | 0.089427 | 0.84369 |
| Denisova | Altai | 7939928 | 46745 | 481452 | 0.089422 | 0.90740 |

Our analysis provides six estimates of coalescence probabilities in the branches of the population tree. If a tree is an accurate representation of the history of these three groups, then these probabilities have to be consistent with one another. The population tree is illustrated in Figure 2, where we denote the common ancestor of the two Neanderthals by *N* and the common ancestor of all three populations by *H*. We distinguish the six estimated values of *c* by the population used to obtain that estimate. Thus, for example, *c*(*AN*;*V*) is the coalescence probability in the branch *AN* using the Vindija genome as population *A*.

**Figure 2.**
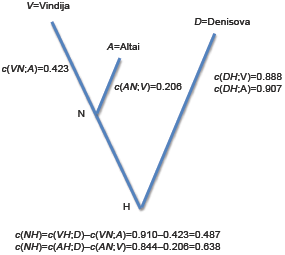
Illustration of the results of applying the method described in the paper for estimating the coalescent probability, *c*, on each of branch of a population tree of three archaic genomes. *c* for the branch leading to Denisovan (DH) can be estimated using either Vindija (V) or Altai (A) as the other population. *c* for the internal branch (NH) can be estimated either by subtracting from either the Vindija or Altai branch. If the history of these populations is accurately represented by a population tree, the estimates of *c* obtained from different subsets of the data should be approximately equal.

There are two tests of consistency with a treelike population history. One is whether the coalescence probability in the branch *DH* is the same when estimated from the Altai and Vindija genomes. As shown on the figure, the two values are similar, 0.888 and 0.907. The second test is whether the same coalescence probability is obtained for the internal branch, *NH*, when estimated two different ways. Because coalescence events in different branches are mutually exclusive, the coalescence probability on the internal branch can be computed by subtracting the value obtained from either the Altai or from the Vindija genomes. As shown, the estimated probabilities are quite different, 0.487 and 0.638, causing us to reject the tree as a correct model. That conclusion is not surprising given that both Prüfer et al. (2014) and Prüfer et al. (2017) concluded that there was gene flow between the Altai and Denisovan populations and admixture into Denisovans from a super-archaic group.

To convert the estimates of *c* to estimates of *T*, we need to solve Equation (5) numerically after assuming something about the history of population sizes in each group. We used the size estimates obtained by Prüfer et al. (2017) from applying PSMC (LI AND DURBIN 2011) to each genome. PSMC returns piecewise constant estimates, with size *N*_*i*_ in time interval (*t*_*i*_,*t*_*i*+1_) with *t*_0_=0. We used the time intervals and sizes reported in Figure S7.5 in Supplement 7 of Prüfer et al. (2017). For piecewise constant population sizes, Equation (5) reduces to

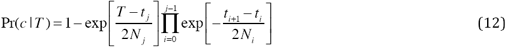

where *j* is chosen so that *t* _*j*_ < *T* ≤ *t* _*j*+1_. Solving Equation (12) yields an estimate of *T* / (2*N*_0_), where *N*0 is different for different populations. For the Vindija and Altai branches, we obtained *T*_*VN*_ / *T*_*AN*_ = 3.041. This ratio is slightly less than ratio of 4 estimated by Prüfer et al. (2017).

## Discussion and Conclusions

We present a simple method to estimate the divergence time of two populations when single genomes are sampled from each population. Our method is a minor modification of a method introduced by Rasmussen et al. (2014). We compare the theoretical basis of our method with two similar methods, one called the *F*(*A* | *B*) method (GREEN *et al.* 2010) that has been used extensively for the analysis of archaic genomes and the other called the TT method (SCHLEBUSCH*et al.* 2017). All three methods are similar in using SNP data from two diploid genomes, one sampled from each of two populations. They all analyze polymorphic SNPs as if they are unlinked. And they all assume a model in which the two populations diverged from one another instantaneously at some time in the past and remained isolated until the genomic samples were taken. None of the methods assumes that the samples are taken at the present and hence they are all applicable to ancient DNA if it is of sufficient quality that heterozygous sites can be called.

The three methods differ slightly in the assumptions they make. The *F*(*A* | *B*) method and the TT method assume that ancestral and derived alleles can be distinguished. Our method does not require that assumption. The *F*(*A* | *B*) method and TT method both require assumptions about the size of the ancestral population. The *F*(*A* | *B*) method assumes a history of population sizes inferred from PSMC (LI AND DURBIN 2011). The TT method assumes that population size was constant before the populations diverged. Our method makes no assumption about the size of the ancestral population. The demography of the ancestral population is captured in the parameter (*k*_1_) that characterizes the folded site-frequency spectrum at the time of population separation.

The three methods differ in which subsets of sites are analyzed. The *F*(*A* | *B*) analyzes all sites that are heterozygous in one population (population *B*) and then computes the fraction of those sites at which a single chromosome from the other population (population A) carries the derived allele. It estimates from this fraction the time of separation from the common ancestor to population *B*. To estimate the divergence time of the other population, A and B are reversed. Our method is similar in focusing on one population at a time, but it analyzes all sites that are polymorphic in an outgroup. The TT method in contrast analyzes all sites polymorphic in the two genomes.

In theory, the three methods differ in how they estimate divergence times. Both the *F*(*A* | *B*) and TT methods estimate the divergence times scaled by the mutation rate. Our method estimates first the coalescence probability in each population and then estimates the coalescence time from some assumption about the history of population sizes after the populations diverged. In practice, the history of population sizes is inferred from PSMC which depends on an assumed mutation rate. Therefore, all three methods depend on the mutation rate. However, the coalescent probabilities estimated with our method and with the TT method do not depend on the assumed mutation rate and hence can be used in the test for a tree-like population history that we have proposed.

One goal of our paper is to call attention to three methods for estimating population divergence times using SNP data from pairs of genomes. These methods have a similar theoretical structure. The differences between them are relatively minor. Most important to the accuracy of results obtained using any of them is the assumption of complete isolation of the populations after they diverged from a common ancestor.

This research was supported in part by NIH Grant GM40282 to M. S. We thank J. Kelso, S. Pääbo and K. Prüfer for helpful discussions of this topic.

## Literature Cited

Green, R. E., J. Krause, A. W. Briggs, T. Maricic, U. Stenzel et al., 2010 A draft sequence of the Neandertal genome. Science 328: 710–722.

Li, H., and R. Durbin, 2011 Inference of human population history from individual whole-genome sequences. Nature 475: 493–496.

Meyer, M., M. Kircher, M.-T. Gansauge, H. Li, F. Racimo et al., 2012 A high-coverage genome sequence from an archaic Denisovan individual. Scienc 338: 222– 226.

Prüfer, K., C. de Filippo, S. Grote, F. Mafessoni, P. Korlević et al., 2017 A high-coverage Neandertal genome from Vindija Cave in Croatia. Science 358: 655.

Prüfer, K., F. Racimo, N. Patterson, F. Jay, S. Sankararaman et al., 2014 The complete genome sequence of a Neanderthal from the Altai Mountains. Nature 505: 43–49.

Rasmussen, M., S. L. Anzick, M. R. Waters, P. Skoglund, M. DeGiorgio et al., 2014 The genome of a Late Pleistocene human from a Clovis burial site in western Montana. Nature 506: 225–229.

Schlebusch, C. M., H. Malmstrím, T. Günther, P. Sjídin, A. Coutinho et al., 2017 Southern African ancient genomes estimate modern human divergence to 350,000 to 260,000 years ago. Science 358: 652.

Schlebusch, C. M., P. Skoglund, P. Sjídin, L. M. Gattepaille, D. Hernandez et al., 2012 Genomic variation in seven Khoe-San groups reveals adaptation and complex African history. Science 338: 374–379.

Skoglund, P., A. Gotherstrom and M. Jakobsson, 2011 Estimation of population divergence times from non-overlapping genomic sequences: examples from dogs and wolves. Molecular Biology and Evolution 28: 1505–1517.

Wakeley, J., 2009 Coalescent Theory. Roberts & Company, Greenwood Village, Colorado.

Zuckerkandl, E., and L. Pauling, 1962 Molecular disease, evolution, and genetic heterogeneity., pp. 189–225 in Horizons in Biochemistry, edited by M. Kasha and B. Pullman. Academic Press, New York.

Zuckerkandl, E., and L. Pauling, 1965 Evolution divergence and convergence in proteins, pp. 97–166 in Evolving Genes and Proteins, edited by V. Bryson and H. J. Vogel. Academic Press, New York.

